# Variation in space and time: a long-term examination of density-dependent dispersal in a woodland rodent

**DOI:** 10.1101/536144

**Authors:** Simon T. Denomme-Brown, Karl Cottenie, J. Bruce Falls, E. Ann Falls, Ronald J. Brooks, Andrew G. McAdam

## Abstract

Dispersal is a fundamental ecological process that can be affected by population density, yet studies report contrasting effects of density on propensity to disperse. Additionally, the relationship between dispersal and density is seldom examined using densities measured at different spatial scales or over extensive time-series. We used 51-years of trapping data to examine how dispersal by wild deer mice (*Peromyscus maniculatus*) was affected by changes in both local and regional population densities. We examined these patterns over both the entire time-series and also in ten-year shifting windows to determine whether the nature and strength of the relationship changed through time. Probability of dispersal decreased with increased local and regional population density, and the negative effect of local density on dispersal was more pronounced in years with low regional densities. Additionally, the strength of negative density-dependent dispersal changed through time, ranging from very strong in some decades to absent in other periods of the study. Finally, while females were less likely to disperse, female dispersal was more density-dependent than male dispersal. Our study shows that the relationship between density and dispersal is not temporally static and that investigations of density-dependent dispersal should consider both local and regional population densities.

## Introduction

Dispersal is defined as movement of an individual from a place of origin to one of reproduction or from one reproductive locale to another (Greenwood 1980), and influences fundamental ecological processes such as population growth rates and viability, genetic connectivity, and range expansions (Greenwood 1980; Nathan 2006; Holyoak et al. 2008). Dispersal is identified as one of four fundamental processes in community ecology (Vellend 2010), and a cornerstone of the metacommunity concept (Leibold et al. 2004). Yet the causes of dispersal are often poorly understood and dispersal remains difficult to measure and include in empirical studies (Koenig et al. 1996; Nathan et al. 2003; Logue et al. 2011).

Benefits of dispersal often stem from differences in conspecific densities between pre and post-dispersal settlement sites. When conspecific densities at pre-dispersal sites are greater than at post-dispersal sites (i.e. positive density-dependent dispersal), dispersing individuals may benefit from reduced competition and agonistic interactions (Murray 1967; Gaines and McClenaghan 1980; Matthysen 2005). Alternatively, negative density-dependent dispersal may be beneficial when individuals disperse to gain access to a limiting resource, or when the benefits of group living outweigh the costs of intraspecific competition (Bowler and Benton 2005).

Conflicting examples of positive and negative density-dependent dispersal (DDD) across a variety of taxa (Bowler and Benton 2005; Matthysen 2005; Rodrigues and Johnstone 2014), have left little consensus regarding general effects of density on dispersal. Matthysen (2005) concluded that both birds and mammals predominantly exhibit positive DDD. The review, however, excluded studies exhibiting temporal trends in density and seasonal variation which Matthysen (2005) acknowledged could have changed their conclusions. Contradicting effects of density on dispersal have also been found within species. DDD has been observed as both negative and positive in separate studies of the same species (Garten and Smith 1974; Rehmeier et al. 2004), or depending on when in the annual cycle density is measured (Betini et al. 2015), or simultaneously negative and positive across a patchily distributed population (Kim et al. 2009).

In particular, variable effects of density on dispersal have been reported in deer mice, *Peromyscus maniculatus*. *Peromyscus* spp. generally display male-biased dispersal (Fairbairn 1978b; Gaines and McClenaghan 1980; Rehmeier et al. 2004) and have been observed to exhibit both positive (Garten and Smith 1974; Fairbairn 1978b; Anderson and Meikle 2010) and negative DDD (Rehmeier et al. 2004; Wojan et al. 2017). These variable effects of density on dispersal suggest that temporal and spatial variation in population densities could feasibly influence the strength of density dependence in a system. Therefore, as others have proposed (e.g. Matthysen 2005), DDD must be examined using long time-series, while accounting for spatial and temporal variation in population densities. In this study, we examined the frequency and extent of dispersal by wild *P. maniculatus*, using monitoring data spanning 50 years from traplines distributed across nearly 17km of temperate forest. These data allowed us to examine dispersal patterns at large temporal and spatial scales and investigate how the strength of DDD was affected by both temporal and spatial variation in population densities.

## Materials and methods

### Data Collection

Deer mice were live-trapped in Algonquin Provincial Park (APP), Ontario, Canada from 1960-2015. Trapping initially occurred on 10 traplines, with lines added and removed intermittently throughout the study such that the same 17 lines were trapped consistently from 1995 – 2015. Lines consisted of 10 trapping stations set along transects at 10m intervals. Early in the study, most lines had one trap at each station, but by 1979 all lines had two traps per station in an attempt to avoid trap saturation, for a total of 20 traps per line. Sherman-style live traps were used exclusively until 2013, when one Longworth trap (Natural Heritage Book Society, Totnes, Devon) replaced one Sherman at each station. Traps were baited with peanut butter and rolled oats until 1991. After 1991, water-soaked sunflower seeds were used as bait. Freeze-killed mealworms (*Tenebrio molitor*) were added to half the traps from 2013-2015 as a bait supplement. Use of Longworth traps and mealworms were aimed at decreasing mortality in shrew (*Soricidae*) species (Do et al. 2013; Shonfield et al. 2013). Distances between lines ranged from 0.115-16.9km. Lines were located in forested habitats with different dominant trees including sugar maple (*Acer saccharum*), white pine (*Pinus strobus*), white spruce (*Picea glauca*), black spruce (*Picea mariana*), trembling aspen (*Populus tremuloides*), and red pine (*Pinus resinosa*).

Each year, trapping was conducted on three consecutive nights either once or twice a month from May through September, for a maximum of 10 three-night trapping sessions per year. Traps were set just before dusk and checked after dawn. When captured, deer mice were weighed, sexed, aged and assessed for reproductive condition. Before release, animals were fitted with uniquely coded ear tags for individual identification upon recapture.

### Dispersal Detection

Dispersal was initially detected when an individual deer mouse was captured on multiple traplines within a single trapping season. Although these data were collected in a consistent manner, any errors in data collection or transfer to digital records could mistakenly appear as dispersal events (e.g. incorrect tag or trapline numbers were recorded). While the error rate is likely low, the compounding of even a low rate of error over more than 50 years of data could skew our estimates of dispersal. To minimize the frequency of such errors we used a strict criterion to determine whether detected movements between traplines represented dispersal events. To be considered a dispersing individual, an individual mouse must have been captured on multiple, consecutive occasions on one line either preceded, or followed by at least one capture on another line. We assumed that the mouse in question was either trapped while moving towards a trapline where it subsequently settled, or was trapped while dispersing from a trapline and either continued on or died. To meet this criterion, a mouse must have been captured on a minimum of three occasions within a year. Accordingly, all model analyses and calculations of dispersal frequencies discussed hereafter were performed on a dataset of mice caught three or more times in a year. Additionally, we avoided including mis-read tags by ensuring that dispersing animals had consistent records of individual traits such as sex and size. Instances where an animal returned to their original line of capture following a capture on a different line were not considered to be dispersal events. These could have represented nomadic wandering between lines rather than dispersal or this could have been caused by a misread tag or data entry error. Animals last caught on their initial line of capture were therefore not considered to have dispersed (n = 136).

Movements occurring between years were not considered as few mice survive the winter in APP, and thus there were relatively few potential dispersers over this period. Movements between lines were considered to be potential dispersal events as the distance between the closest traplines (115m) was greater than the diameter of home ranges for this species typically reported in the literature (Stickel 1968; Bowers and Smith; 1979; Harestad and Bunnel 1979; Van Horne 1980; Abramson et al 2006; Wood 2010). Distances between traplines were calculated as straight-line distance between the first trap station on each line. Finally, changes in methodology and some loss of data necessitated the exclusion of five years of data from the dataset (1987-1991).

### Density Measures

We converted raw capture totals to the number of captures per hundred trap-nights per year to account for variation in trapping effort both between years and among traplines. The first measurement was raw local trapline density (hereafter termed raw local density. We converted these raw local densities into two types of densities for the statistical analyses that determined the independent effect of regional and local density on dispersal. The first density measure was the average regional trapline density, calculated annually as the mean number of captures per hundred trap-nights across all traplines in APP in that year (or regional density) Second, we calculated relative local density (hereafter local density) for each trapline in each year by subtracting the regional density in a year from the raw local density on each line that same year. This local density measure thus captures the additional effect of local densities after accounting for year-to-year fluctuations in regional desntiy associated with for instance seed mast events (Falls et al. 2007). This relative local density thus allowed us to include uncorrelated local densities and regional density estimates within the same models (*sensu* van de Pol and Wright 2009).

It should be noted that the density estimates we used do not account for spatial or temporal variation in capture probability. While spatially explicit mark-recapture techniques (SECR) represent a powerful set of statistical tools to account for these issues (Royle et al. 2013), they were inappropriate for our long-term data for several reasons. First, Bayesian SECR models are inappropriate for trapping data that are derived from single-catch traps such as those used in our study (Gerber and Parmenter 2015). Second, when using frequentist SECR models the multi-catch estimator can only be used if density is relatively constant over survey regions (Distiller and Borchers 2015), and this was not the case in our data (see Results). Finally, the single-catch estimator developed by Distiller and Borchers (2015), which allows for frequentist SECR analysis using single catch traps, requires times for all captures which were not available for our historical data. We discuss this issue further below.

### Data Analysis

To assess how a deer mouse’s propensity to disperse was affected by sex and population density at different spatial scales, we fitted eight separate Generalized Linear Mixed Effects Models (binomial family, logit link) associated with local and regional population densities as well as sex of the individual mouse. They were 1) Biological Null Model, 2) Sex Model, 3) Local Density Model, 4) Regional Density Model, 5) Raw Density Model (RWD), 6) Local and Regional Density Model, 7) Density Interaction Model (DI), 8) Density and Sex Interaction Model (DSI) (see Table 1). All eight models used the binary response variable of whether an individual mouse dispersed or not and included the fixed effect of distance (in meters) from the trapline where the mouse was initially caught to the next closest trapline. This effect was included to account for the non-uniform spatial distribution of traplines and whether differences in movement among lines were due primarily to spatial effects on the probability of detecting a dispersal event (i.e. dispersal is more likely to be detected between nearby traplines). We used the natural log of the raw distance in kilometers (km) to correct a significant non-linearity in the relationship between this variable and the probability of dispersal. In all models, local density and regional density were converted to z-scores 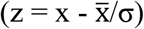 based on the mean and standard deviation of all observations across years and lines.

**Table 1:** Eight generalized linear mixed effects models (binomial response, logit link) fitted to the data for predicting the probability of an individual deer mouse dispersing

| Model | K | AIC | ΔAIC | LogLik | AIC $\omega$ | Rank |
| --- | --- | --- | --- | --- | --- | --- |
| Density and sex interaction model (DSI): ∼ Local Density + Regional Density<sup>+</sup> + Sex<sup>‡</sup> + Closest Distance<sup>¶</sup> + Local Density:Sex + Local Density:Regional Density | 9 | 1059.8 | 0 | -520.8 | 0.686 | 1 |
| Density interaction model (DI): ∼ Local Density + Regional Density + Sex + Closest Distance + Local Density:Regional Density | 8 | 1062.0 | 2.2 | -523.0 | 0.228 | 2 |
| Local and regional density model : ∼ Local Density + Regional Density + Sex + Closest Distance | 7 | 1065.3 | 5.5 | -525.7 | 0.044 | 4 |
| Raw density model (RWD): ∼ Raw Density + Sex + Closest Distance | 6 | 1065.6 | 5.7 | -526.8 | 0.040 | 3 |
| Regional density model: ∼ Regional Density + Sex + Closest Distance | 6 | 1071.4 | 11.6 | -529.7 | 0.002 | 5 |
| Local density model: ∼ Local Density + Sex + Closest Distance | 6 | 1074.7 | 14.9 | -531.4 | <0.001 | 6 |
| Sex model: ∼ Sex + Closest Distance | 5 | 1081.0 | 21.2 | -535.5 | <0.001 | 7 |
| Biological null model: ∼ Closest Distance | 4 | 1085.4 | 25.6 | -538.7 | <0.001 | 8 |

Note: K = number of parameters, AIC = Akaike Information Criterion, ΔAIC = Model's AIC -the lowest AIC of any model, LogLik = log likelihood, AIC $\omega$ = Aikaike Weight.
Random effects in both models were year and trapline of emigration.
\* density (captures/hundred trap-nights) of deer mice on the trapline an individual is initially captured on minus the regional density in that year
<sup>+</sup> density (captures/hundred trap-nights) of deer mice averaged across all traplines in a year
<sup>‡</sup> whether an animal is male or female, male used as reference category
<sup>¶</sup> natural log of the distance in metres from the line of an individual's initial capture to the next most closely situated line

We used the Akaike Information Criterion (AIC) to assess relative support for the eight models (Burnham and Anderson 2007), which included various combinations of sex, and raw, local and regional density as fixed effects (Table 1). We calculated AIC values for each model and performed model comparisons using ΔAIC (Δ_i_) values (Table 1). Akaike Weights (ω_i_) were calculated to examine the conditional probabilities of each model (Table 1). All models contained the same random effects, with initial trapline of capture and year of capture included to capture spatial and temporal variation in dispersal propensity. Likelihood ratio tests (LRT) were used to assess the significance of the year and trapline random effects in each of the 8 candidate models. Models were assessed for overdispersion and collinearity of variables and met model assumptions. Statistical analyses were performed in R (Version 3.5; R Core Team 2018), including the glmer function with a bobyqa optimizer in the lme4 package (Bates et al. 2015).

In addition to the model analysis described above, we performed a number of *post hoc* analyses. When first visualizing the relationship between regional density and movement frequency (Figure 1a) there appeared to be a marked change in the relationship between the two variables for portions of the time-series, particularly over a 9-year period from 2002-2010. To examine this, we ran model with raw local density as a fixed effect and *year* and *trapline* of initial capture as random effects on a shifting ten-year window through the time-series. This model was used because the RWD model was the simplest model with moderate support based on AIC comparisons and contained only raw local density and sex as fixed effects. More complex models would not fit these data given the smaller sample size provided in 10-year windows of the time-series (Figure 1b).

**Fig 1:**
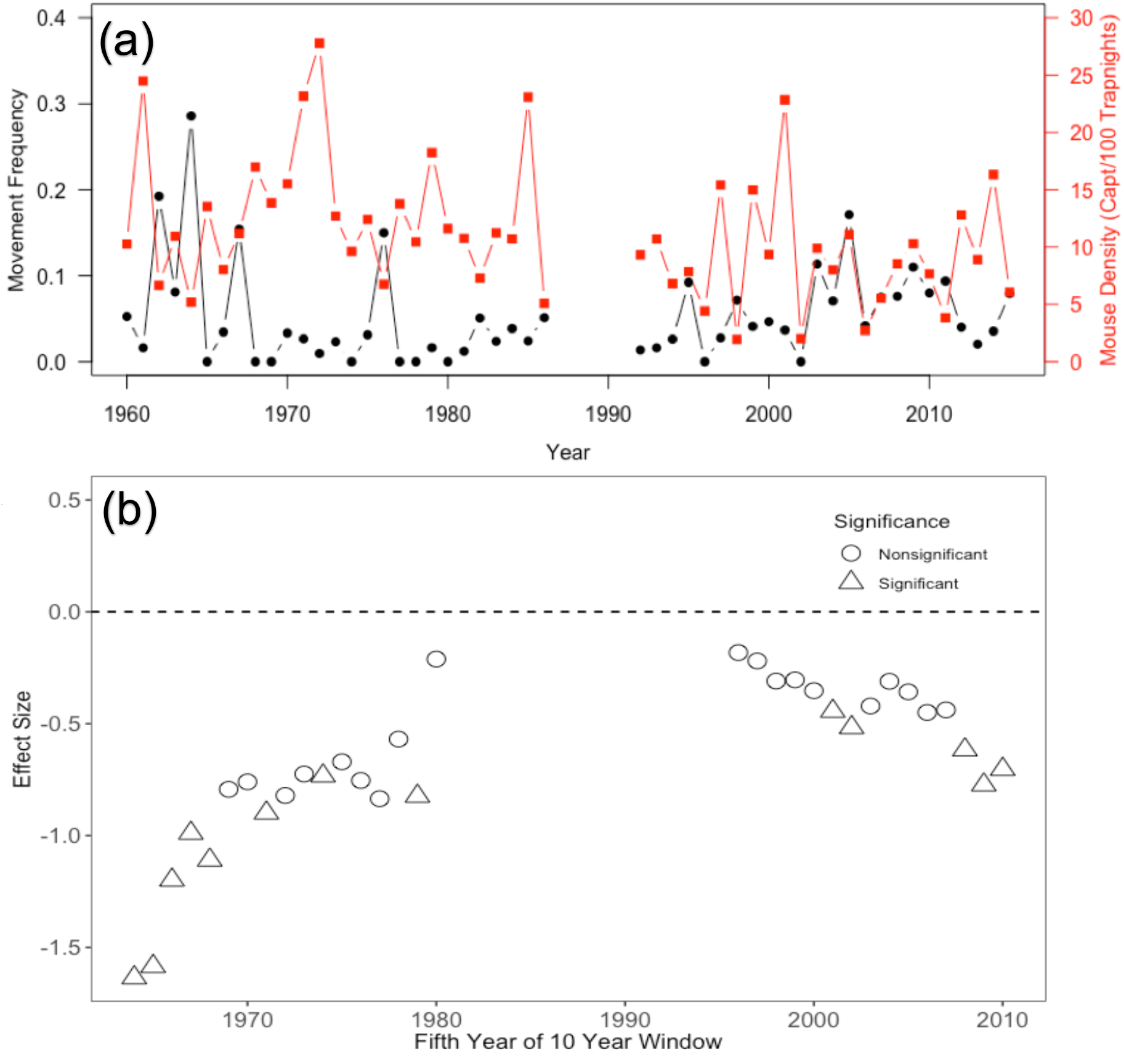
a) Proportion of possible dispersers that dispersed each year plotted alongside the density of the deer mouse (*Peromyscus maniculatus*) population in APP between 1960-2015. b) the effect size and significance of the relationship between raw density and dispersal probability as analyzed in 32 ten-year shifting windows using a model with raw local density as a fixed effect and *year* and *trapline* of initial capture as random effects. The magnitude of this effect of raw density on the probability of dispersing (effect size) is plotted against the midpoint of the 10-year window. Windows where the relationship is significant are represented with triangles and lack of significance is represented with squares. Significance determined at ⍺ = 0.05.

The second set of *post hoc* analyses assessed how robust our results were to changes in trapping protocols during this long-term study (e.g. number of traps per station, change in bait, and trap types), that occurred at distinct points in our study (1979, 1991 and 2013 respectively). For this we ran our best model on subsets of our dataset from before, between and after these protocol changes occurred.

## Results

### Population Densities and Frequency of Movement

Over the 51 years included in this study, there were 3408 instances where a mouse was caught at least three times in a year. Of these 3408 possible dispersers, 4.2% (n = 142) dispersed between traplines. The proportion of the population that dispersed fluctuated between years (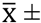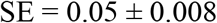 mice dispersing in the population, n = 51 years), with no dispersal detected in some years and a maximum of 28.6% of the population dispersing between lines in 1964 (Figure 1). The population sex ratio was slightly skewed, with males comprising 53.4% of individuals in the dataset (X^2^ = 7.9, df = 1, P < 0.01). This male skew was more pronounced among dispersing individuals, with 64.8% of dispersing mice being male (X^2^ = 7.7, df = 1, P < 0.01). Overall, males were more likely to disperse (5.1%) than females (2.7%; Fisher Exact Test: P = 0.006).

Distances travelled by dispersing individuals ranged from 115m to 11.4 km 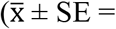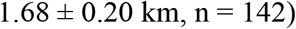 (Figure 2). There was no difference in dispersal distance between the sexes (t = 0.98, df = 92, p-value = 0.3) (Figure 2). Only 51.4% (n = 73) of dispersals were to the next most closely situated trapline (Figure 2). There were two traplines located only 115m apart, but dispersal between these traplines represented only 16.9% (n = 24) of all movements.

**Fig 2:**
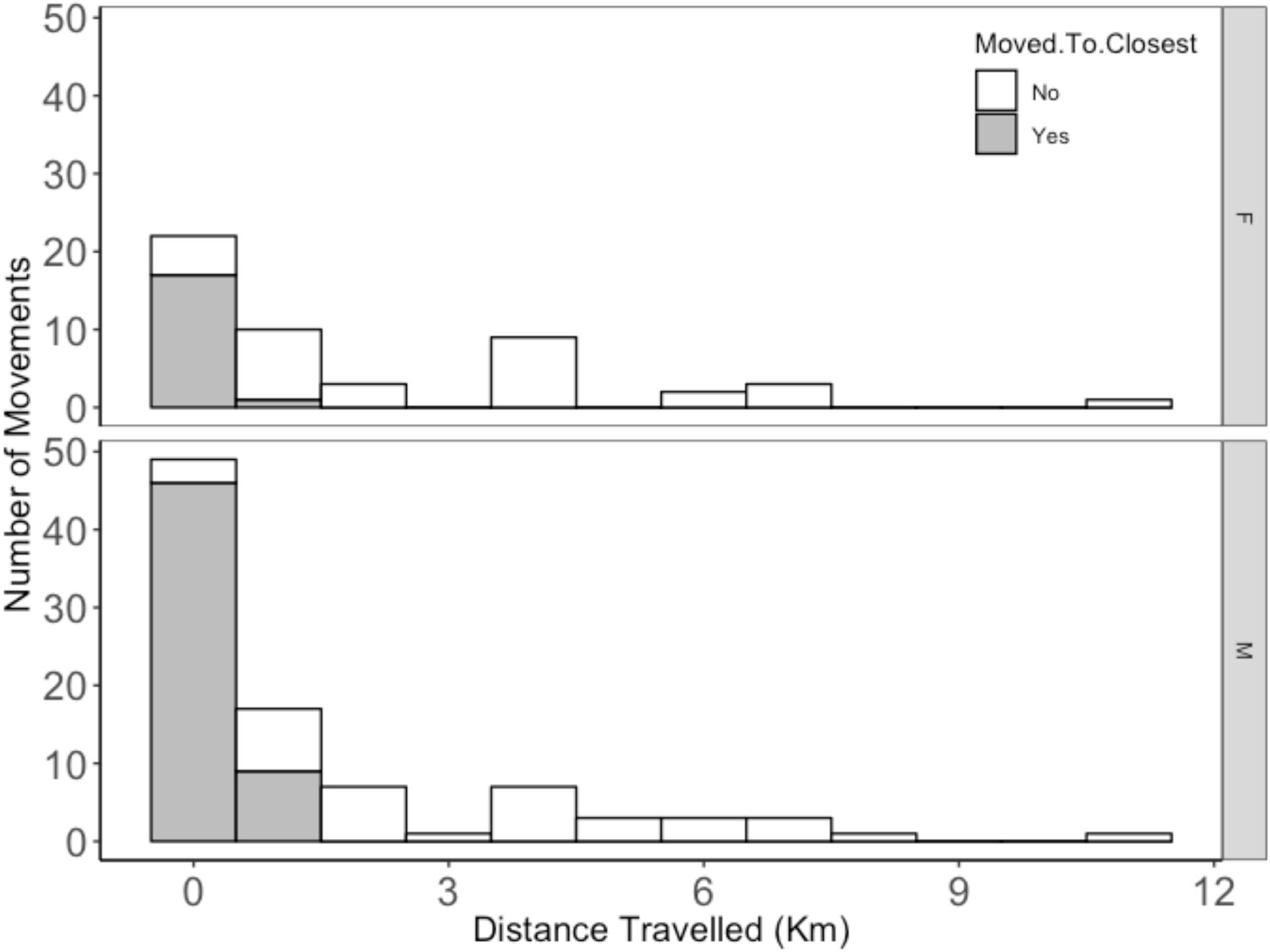
Distances travelled by dispersing deer mice (N = 142). (F) is for females while (M) represents males. The number of individuals that dispersed to the closest trapline is depicted by the height of the open bars, whereas those that dispersed farther than the nearest trapline are depicted by the height of the shaded bar. There was no difference in dispersal distance between the sexes (t = 0.98, df = 92, p-value = 0.3)

The density of deer mice in APP fluctuated over an order of magnitude during this study, with yearly regional population densities ranging from 1.9 to 27.8 captures per hundred trap nights (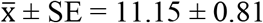 captures per hundred trap-nights, n = 51 years) (Figure 1a). Local densities also varied, with the average range in local population densities among traplines within years (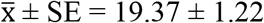 captures per hundred trap-nights, n = 51 years) approaching the aforementioned range in regional densities among years. In particular, 9 of 51 years exhibited a greater range in density among lines within a year than the range in regional density across the 51 years, with a maximum range of 47.2 captures per hundred trap-nights in 2001.

### Dispersal Models

The model that had the lowest AIC score of the eight fitted models was the DSI model which included the fixed effects of both regional and local density, sex, and interactions between both local density and regional density, and local density and sex (Table 2). In the DSI model, the probability of dispersal decreased with increasing distance to nearest trapline (β ± SE = −1.035 ± 0.15, Z = −6.85, P < 0.0001), increased regional density (β ± SE = −0.63 ± 0.16, Z = −3.9, P < 0.0001), and increased local density (β ± SE = −0.091 ± 0.026, Z −3.52, P = 0.0004). The significant positive interaction between local and regional densities (β ± SE = 0.036 ± 0.015, Z = 2.44, P = 0.015) indicated that in years of low regional density, local densities had a stronger negative effect on the probability of dispersal (Figure 3a). For example, three of the five years displaying the highest proportions of dispersers in the study were 1964 (0.286), 1962 (0.192) and 1976 (0.150), which represented three of the 11 lowest density years within the study.

**Table 2:** Summary of the statistical results from density and sex interaction model (DSI)

| Variable – Variable Type | $\beta \pm SE$ | Z | $\partial^2$ | X<sup>2</sup> | P |
| --- | --- | --- | --- | --- | --- |
| Closest Distance - Fixed | -1.035 ± 0.15 | -6.85 | - | - | <0.0001 |
| Sex (Male) - Fixed | 0.32 ± 0.19 | 1.69 | - | - | 0.09 |
| Local Density - Fixed | -0.091 ± 0.026 | -3.52 | - | - | 0.0004 |
| Regional Density - Fixed | -0.63 ± 0.16 | -3.98 | - | - | <0.0001 |
| Local Density : Reg. Density - Fixed | 0.036 ± 0.015 | 2.44 | - | - | 0.01 |
| Local Density : Sex (male) - Fixed | 0.064 ± 0.031 | 2.10 | - | - | 0.04 |
| Year - Random | - | - | 1042 | 0.2 | 0.65 |
| Initial Trapline – Random | - | - | 1042 | 11.2 | 0.0008 |

Note: This model best fitted the observed dispersal data and had strong support based on model comparisons using AIC.
Likelihood ratio tests were used to assess the significance of each random effect.

**Fig 3:**
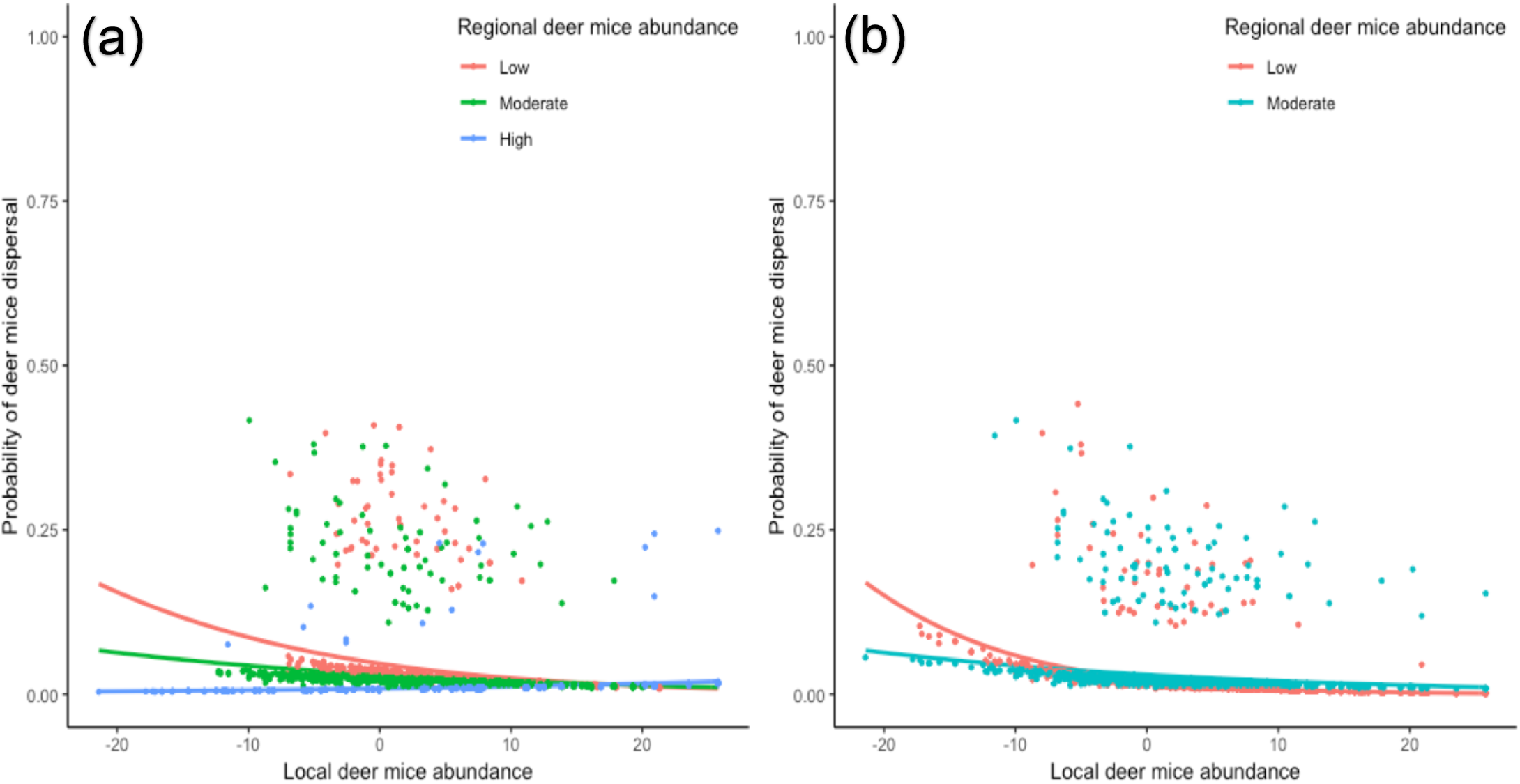
a) Partial plot of the interaction between Regional Density and Local Density from the DSI model (β ± SE = 0.036 ± 0.015, Z = 2.44, P = 0.015). Regional Density is grouped by tertile. The positive interaction can be seen to make the negative density-dependent dispersal in response to local density more pronounced when regional density was low (green trend-line). When regional density was high (pink trend-line) there was no effect of local density. b) Partial plot of the interaction between Local Density and Sex (F=Female, M = Male) from the DSI model (β ± SE = 0.064 ± 0.031, Z = 2.10, P = 0.036). While males are more likely to disperse, this interaction is approaching significance and suggests that female dispersal may be more sensitive to changes in local density than male dispersal.

The DSI model also included sex as fixed effect and while more males dispersed than females, the effect of sex on dispersal was not significant (β ± SE = 0.32 ± 0.19, Z = 1.69, P = 0.09) (Figure 3b). However, the interaction between local density and sex was in fact significant (β ± SE = 0.064 ± 0.031, Z = 2.1, P = 0.036), indicating that while males disperse more than females, females were more sensitive to local density than males and were more likely to disperse when densities were low (Figure 3b). Initial *line of capture* was a significant random effect in this model (LRT, Deviance = 1042, X^2^_1_= 11.2, P = 0.0008) whereas *year* was not (LRT, Deviance = 1042, X^2^_1_ = 0.2, P = 0.65). Overall, the DSI model explained 11.77% of the deviance.

There was nearly strong support (Δ_i_ < 2) for the DI model(Δ_i_ = 2.2) which was identical to the DSI model except it did not include an interaction between local density and sex. All effects in this model were in the same direction as in the DSI model and the DI model explained 10.05% of the overall deviance. Considerably less support was seen for models that did not contain the interaction between local and regional densities, with all these models demonstrating either moderate (Δ_i_ < 7) or no support (Δ_i_ > 10) (Table 1).

Despite evidence of negative DDD overall, visual inspection of the time series of density and dispersal probability (Figure 1a) suggested temporal changes in the strength of density-dependent dispersal. When running the simplified model containing raw local density as a fixed effect and *year and trapline* of initial capture as random effects on the shifting 10-year window through the time series, the effect of raw density on dispersal probability never became positive. The effect size did approach zero, however, and the negative relationship between raw density and dispersal probability was significant for only 13 of 32 ten-year windows in the analysis (Figure 1b). Finally, when running the DSI model on our data before and after each change in trapping methodology, there were no changes in the direction of effects for all explanatory variables save one. The exception was the effect of sex on dispersal, which was reversed prior to the initial bait change, but the effect was not significant.

## Discussion

### Probability of Dispersal

Overall, deer mice in APP exhibited negative DDD, with a lower probability of dispersal at high densities, regardless of the scale at which density was measured. Proposed mechanisms for negative DDD include attraction to conspecifics (Danielson and Gaines 1987; Stamps 1991), range constriction in the face of high densities (Stickel 1960; Taitt 1981) and exclusionary behavior towards immigrants in periods of high density (i.e. “social fences”; Hestbeck 1982). Had trap saturation occurred in APP it could mimic a natural social fence, denying immigrants access to traps on densely populated lines and biasing our results towards the appearance of negative DDD. This seems unlikely, however, as densities on even the most highly populated traplines fell well short of saturation (i.e. average captures per 100 trap-nights across all lines throughout this study was 22.62 with SE = 0.42 and SD = 10.66; the highest capture rate on any line in any year was 73.81 captures per 100 trap-nights). Additionally, there was no obvious change in the direction of density-dependence after the traps per station were doubled in 1979.

While the result of negative DDD runs counter to generalizations of positive DDD in mammals (Matthysen 2005), previously reported relationships between dispersal and density in *Peromyscus* spp. are inconsistent. Positive DDD has been observed in *P. maniculatus* (Fairbairn, 1978a), *P*. *polionotus* (Garten and Smith 1974), and genetic estimates show *P. leucopus* is more likely to disperse from high density habitat patches (Anderson and Meikle 2010). Alternatively, there is evidence of negative DDD in both *P*. *maniculatus* (Rehmeier et al. 2004) and *P*. *boylii* (Wojan et al. 2015). The spatial and temporal breadth of our study may elucidate why studies on *Peromyscus* spp., and other dispersing organisms, often yield divergent results. For while deer mice in our study consistently exhibited negative DDD, the strength with which density affected dispersal varied temporally, spatially and between sexes.

The relationship between density and dispersal varied throughout our time-series. This variation is evident when examining the time-series of regional density and movement frequency together (Figure 1a), particularly the period from 2002-2010 which visually suggests a possible positive relationship. The effect of density on dispersal probability in the RWD model was never positive in any 10-year window along the time-series, however, the effect size approached zero in the 1990’s before becoming increasingly negative (Figure 1b). Also, the majority of the 10-year windows examined (n = 19) did not exhibit significant relationships between density and dispersal. Few studies investigating DDD have used long-term data (Matthysen 2005), despite the fact that inferences drawn from short-term studies can differ drastically from those generated by long-term research (Kratz et al. 2003), and that effects of some ecological processes become more pronounced over time (Cardinale et al. 2007). In an experiment using guppies (*Poecilia reticulata*), De Bona et al. (2019) recently demonstrated that over the course of a colonization event, density-dependent dispersal can change in strength and direction. Still, our findings are the first to demonstrate that the density-dispersal relationship can vary through time in an unmanipulated system and over large time-series. This highlights the value of long-term studies for drawing accurate conclusions regarding the density-dispersal relationship, as an unfortunately timed relatively long-term study (i.e. 10 years) would not detect the negative effect of density on dispersal that exists in this system.

In addition to varying temporally, the strength with which local density affected dispersal was mediated by regional densities. This interaction between the regional and local density effects on dispersal indicates that the negative effect of local density on dispersal is greater in years of low regional density and muted in years of high regional density (Figure 3a). This suggests that local density can be an important driver of dispersal, but only when examined in concert with regional density, as low local densities that increase the probability of dispersal are only noteworthy in years when regional densities are also low. Local and regional densities acting as interacting drivers of dispersal has been observed experimentally (De Bona et al. 2019) but never before in natural, unmanipulated systems. Considering local densities within the context of the accompanying regional densities is important moving forward, as many studies provide only densities for localized sites of emigration (Matthysen 2005), which based on our findings may lead to spurious results when not considered in the context of associated regional densities. As well, the few large datasets that have been used to examine DDD fail to differentiate between spatial and temporal variation in densities (Matthysen, 2005; but see Pasinelli and Walters 2002; Catchpole et al. 2004) despite the fact both temporal variation and interactions between densities at multiple spatial scales affect dispersal.

Several potential mechanisms could cause this interaction between the effects of local and regional densities on dispersal. Deer mice populations in APP are closely tied to maple seed crops, with mouse populations peaking after seed masting (Falls et al. 2007). *Peromyscus* spp. also move based on habitat quality and population density (Morris and Diffendorfer 2004). It may be that individuals select habitat and disperse to high quality, maple-rich areas when possible; but in years of high regional densities, local densities in high-quality habitats rise enough to prevent such immigration. This process would strike a balance between traditional dispersal hypotheses predicting positive DDD due to increased competition (Greenwood 1980), and ideas of conspecific attraction (Stamps 1991) and habitat selection. This balance between competition and facilitation amongst conspecifics has been observed previously in blue-footed booby (*Sula nebouxii*), where simultaneous positive and negative DDD results from the costs of competition and the advantages of increased mating choice options (Kim et al. 2009).

Mate availability may also influence dispersal in APP as the strength of density-dependence also differed between sexes. Males dispersed more often than females, which is consistent with findings of male-biased dispersal in both *Peromyscus* spp. (Fairbairn 1978a; Goundie and Vessey 1986; Rehmeier et al. 2004; Mabry 2014), and other mammals (Greenwood 1980; Dobson 1982; Mabry et al. 2013). However, we also found evidence that dispersal in females was more sensitive to changes in local density than in males, with females dispersing more frequently than males when local densities were very low. Examples of sex-biased dispersal switching from male-bias to female-bias within a system are rare (but see Pérez-González and Carranza 2009), but have implications for understanding how mating strategies influence dispersal. The predominance of male-biased dispersal in mammals (and female-bias in birds) has long been attributed to mating systems, although the simplicity of this relationship has been questioned (Mabry et al. 2013). If males disperse to track females and females rely on available resources for reproduction (Emlen and Oring 1977), then at exceedingly low densities the lack of available mates may outweigh the benefits of familiar surroundings, resulting in increased female dispersal. Alternatively, if low densities indicate low food availability, this would affect female dispersal to a greater extent due to their reliance on resources for reproduction (Emlen and Oring 1977). Differences between male and female dispersal sensitivity to local densities mean that spatial variation in density could potentially affect sex ratios, in turn affecting population density.

As mentioned previously, our density measures did not account for variation in capture probabilities. While we described the practical reasons for not being able to utilize SECR methods, we recognize that this is not ideal. However, the lack of trap saturation on any of our lines throughout the study, the fact that frequency of recapture of individuals within this study 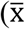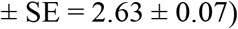 did not decrease with increased regional annual densities (linear regression; β ± SE = 0.012 ± 0.01, t = 1.09, P = 0.28), and the fact that the density-dispersal relationships in our models were not altered by the changes in trapping methodologies that occurred throughout this study all speak to the robustness of our findings.

### Frequency and Distance of Dispersal

Generally, deer mice dispersal occurred as often as expected based on existing literature, but to greater distances. Rehmeier et al. (2004) reported that over a 10-year period 4% of deer mice in a tallgrass prairie undertook long-distance dispersals (LDDs, >125m); Bowman et al. (2001b) similarly reported 4.2% of deer mice demonstrating LDD (>100m) in temperate forests. Based on these papers, our dispersals (>115m) can be classified as LDDs and the frequency of LDD by deer mice in APP (4.2%) was remarkably similar to these frequencies reported in the literature. The dispersal distances in our study exceeded previous reports for *P. maniculatus*, with 13% (n =20) of observed dispersals exceeding the previous reported maximum long-distance movement (4287 ± 10m, in Wood and McKinney 2015). Our observed maximum dispersal distance of 11.3km represents a 164% increase on that previous maximum. As well, we detected more frequent movements of great distances than previous studies, with 3.1% of mice dispersing over 200m compared to 1% of mice dispersing beyond 200m in previous studies (Brant 1962; Groves and Keller 1986). We also observed 2.7% and 2.1% of mice dispersing beyond 300m and 500m respectively, while roughly 1% of populations travelled those distances in previous studies (Diffendorfer et al. 1999; Rehmeier et al. 2004). This indicates that the tail of the dispersal distance distribution of this species has previously been underestimated.

Dispersal distances tend to follow leptokurtic distributions; many individuals disperse short distances with few LDDs. Understanding the rate and extent of LDDs, represented by the tail of the distribution of dispersal distances, is important as they determine the spread of a species in a landscape (Kot et al. 1996; Levin et al. 2003; Green and Figuerola 2005). Species with “fat-tailed” dispersal distributions exhibit more rapid spread throughout landscapes (Kot et al. 1996). This can influence persistence in the face of climate change (Schloss et al. 2012), which small mammals in this region are already responding to (Myers et al. 2009), and the spread of zoonotic diseases (Mundt et al. 2009), of which *Peromyscus* spp. are vectors (Meerburg et al. 2009; Gaitan and Millien 2016). These distribution tails are notoriously difficult to quantify (Koenig et al. 1996), and LDD in deer mice is generally reported anecdotally or with little detail (Jung et al. 2005; Wood and McKinney 2015). Our absolute frequencies of dispersal should be interpreted cautiously as mark-recapture studies have limited ability to detect dispersal. The majority of small mammal studies utilize trapping grids too small to detect LDDs (< 2ha, from Bowman et al. 2001a), and comparisons with radio-tracking and genetic dispersal estimates show that mark-recapture methods underestimate dispersal distances (Koenig et al. 1996). Unfortunately, expanding trapping grids is often impractical, as increasing the maximum distance across a grid exponentially increases area and monitoring effort. Our study takes an alternate approach by instead using many smaller transects, reducing our ability to detect individual dispersal events, but overcoming this via extensive temporal replication. Our results suggest that deer mice LDDs are more common than the literature suggests, reinforcing claims that relatively small-scale mark-recapture monitoring techniques fail to accurately capture LDD (Koenig et al. 1996).

We have demonstrated that deer mice exhibit negative DDD that is more pronounced during periods of low regional densities and that female dispersal is more sensitive to these changes in density than that of males. We also have demonstrated that the relationship between density and dispersal is not static but rather varies conspicuously over large time scales. These results show that long-term field studies are beneficial to the examination of ecological processes such dispersal, as smaller scale projects are likely to miss or potentially misrepresent the broader signal entirely. As well, our findings suggest that while local population densities are important, their influence on dispersal can be affected by broader regional population densities. As such, future studies should account for both local and regional densities when testing for DDD. Additionally, we have shown that well-established phenomena such as mammalian male-biased dispersal may also be density-dependent. These results together suggest that the conflicting relationships between density and dispersal in the literature may result from broader context-dependence in DDD. Finally, our results add to a previously modest body of empirical literature examining LDD, by demonstrating that LDD by deer mice is not uncommon. This reinforces that dispersal distributions estimated from short-term mark recapture studies may be underestimated, which is potentially important for future conservation.

## Acknowledgements

We would like to acknowledge countless individuals who collected these data. Thank you to the Algonquin Wildlife Research Station for logistical support. This work was supported through funding from the National Science and Engineering Research Council of Canada and the Ontario Ministry of Natural Resources. We dedicate this work to the memory of our colleague Dr. Leslie Rye who said it best: “I can’t believe I am getting paid to work in Algonquin!”

## References

Abramson G, Giuggioli L, Kenkre VM, Dragoo JW, Parmenter RR, Parmenter CR, Yates TL (2006) Diffusion and home range parameters for rodents: Peromyscus maniculatus in New Mexico. Ecol Complex 3: 64–70 doi: 10.1016/j.ecocom.2005.07.001

Anderson CS, Meikle DB (2010) Genetic estimates of immigration and emigration rates in relation to population density and forest patch area in Peromyscus leucopus. Cons Gen 11:1593–1605 doi: 10.1007/s10592-009-0033-8

Bates D, Maechler M, Bolker B, Walker S (2015) Fitting linear mixed-effects models using lme4. J Stat Soft 67:1–48 doi: 10.18637/jss.v067.i01

Betini GS, Pardy A, Griswold CK, Norris DR (2015) The role of seasonality and non-lethal carry-over effects on density-dependent dispersal. Ecosphere 6:art272 doi: 10.1890/ES15-00257.1

Bowers MA, Smith HD (1979) Differential habitat utilization by sexes of deermouse, Peromyscus maniculatus. Ecology 60:869–875 doi: 10.2307/1936854

Bowler DE, Benton TG (2005) Causes and consequences of animal dispersal strategies: relating individual behaviour to spatial dynamics. Biol Rev Camb Philos Soc 80:205–225 doi: 10.1017/S1464793104006645

Bowman J, Corkum CV, Forbes GJ (2001a) Spatial scales of trapping in small-mammal research. Can Field-Nat 115:472–475

Bowman J, Forbes G, Dilworth T (2001b) Distances moved by small woodland rodents within large trapping grids. Can Field-Nat 115:64–67

Brant DH (1962) Measures of the movements and population densities of small rodents. Univ Calif publ zool 62:105–184

Burnham KP, Anderson DR (2007) Model selection and inference: A practical information-theoretic approach, 2nd edn. Springer, New York, NY

Cardinale BJ, Wright JP, Cadotte MW, Carroll IT, Hector A, Srivastava DS, Loreau M, Weis JJ (2007). Impacts of plant diversity on biomass production increase through time because of species complementarity. Proc Natl Acad Sci 104:18123–18128 doi: 10.1073/pnas.0709069104

Catchpole EA, Fan Y, Morgan BJT, Coulson TN, Clutton-Brock TH (2004). Sexual dimorphism, survival and dispersal in red deer. J Agric Biol Environ Stat 9:1–26 doi: 10.1198/1085711043172

Danielson BJ, Gaines MS (1987) The influences of conspecific and heterospecific residents on colonization. Ecology 68:1778–1784 doi: 10.2307/1939869

De Bona S, Bruneaux M, Lee AEG, Reznick DN, Bentzen P, López-Sepulcre A (2019) Spatio-temporal dynamics of density-dependent dispersal during a population colonisation. Ecol Lett doi: 10.1111/ele.13205

Diffendorfer JE, Gaines MS, Holt RD (1999) Patterns and impacts of movements at different scales in small mammals. In: Barrett GW, Peles JD (eds) Landscape ecology of small mammals. Springer-Verlag, New York, pp 63–88

Distiller G, Borchers DL (2015) A spatially explicit capture-recapture estimator for single-catch traps. Ecol Evol 21:5074–5087 doi: 10.1002/ece3.1748

Do R, Shonfield J, McAdam AG (2013) Reducing accidental shrew mortality associated with small-mammal livetrapping II: a field experiment with bait supplementation. J Mammal 94:754–760 doi: 10.1644/12-MAMM-A-242.1

Dobson FS (1982) Competition for mates and predominant juvenile male dispersal in mammals. Anim Behav 30:1183–1192 doi: 10.1016/S0003-3472(82)80209-1

Emlen ST, Oring LW (1977) Ecology, sexual selection and the evolution of mating systems. Science 197:215–223 doi: 10.1126/science.327542

Fairbairn, DJ (1978a). Dispersal of deer mice, Peromyscus maniculatus - Proximal Causes and Effects on Fitness. Oecologia 193:171–193 doi: 10.1007/BF00366070.

Fairbairn DJ (1978b) Behaviour of dispersing deer mice (Peromyscus maniculatus). Behav Ecol Sociobiol 3:265–282 doi: 10.1007/BF00296313

Falls JB, Falls EA, Fryxell JM (2007) Fluctuations of deer mice in Ontario in relation to seed crops. Ecol Monogr 77:19–32 doi: 10.1890/05-1485

Gaines MS, McClenaghan LR (1980) Dispersal in small mammals. Annu Rev Ecol Syst 11:163–196 doi:10.1146/annurev.es.11.11080.001115

Gaitan J, Millien V (2016) Stress level, parasite load, and movement pattern in a small-mammal reservoir host for Lyme disease. Can J Zool 94:565–573 doi: 10.1139/cjz-2015-0225

Garten CT, Smith MH (1974) Movement by oldfield mice and population regulation. Acta Theriol 19:513–514

Gerber BD, Parmenter RR (2015) Spatial capture-recapture model performance with known small-mammal densities. Ecol Appl 25:695–705 doi: 10.1890/14-0960.1

Goundie TR, Vessey SH (1986) Survival and dispersal of young white-footed mice born in nest boxes. J Mammal 67:53–60 doi: 10.2307/1381001

Green AJ, Figuerola J (2005) Recent advances in the study of long-distance dispersal of aquatic invertebrates via birds. Divers Distrib 11:149–156 doi: 10.1111/j.1366-9516.2005.00147.x

Greenwood PJ (1980) Mating systems, philopatry and dispersal in birds and mammals. Anim Behav 28:1140–1162 doi:10.1016/S0003-3472(80)80103-5

Groves CR, Keller BL (1986) Ecological characteristics of small mammals on a radioactive waste disposal area in southeastern Idaho. Am Midl Nat 46:404–410 doi: 10.2307/2425405

Harestad AS, Bunnel FL (1979) Home range and body weight --a reevaluation. Ecology 60:389–402 doi: 10.2307/1937667

Hestbeck JB (1982) Population regulation of cyclic mammals: the Social Fence Hypothesis. Oikos 39:157–163 doi: 10.2307/3544480

Holyoak MR, Casagrandi R, Nathan R, Revilla E, Spiegel O (2008) Trends and missing parts in the study of movement ecology. Proc Natl Acad Sci USA 105:19060–19065 doi: 10.1073/pnas.0800483105

Jung T, O’Donovan K, Powell T (2005) Long-distance movement of a dispersing deer mouse, Peromyscus maniculatus, in the boreal forest. Can Field-Nat 119:451–452 doi: 10.22621/cfn.v119i3.161

Kim SY, Torres R, Drummond H (2009) Simultaneous positive and negative density-dependent dispersal in a colonial bird species. Ecology 90:230–239 doi: 10.1890/08-0133.1

Koenig W, Van Vuren D, Hooge PN (1996) Detectability, philopatry and the distribution of dispersal distances in vertebrates. Trends Ecol Evol 11:514–517 doi: 10.1016/S0169-5347(96)20074-6

Kot M, Lewis MA, Van Den Driessche P (1996) Dispersal data and the spread of invading organisms. Ecology 77:2027–2042 doi: 10.2307/2265698

Kratz TK, Deegan LA, Harmon ME, Lauenroth WK (2003) Ecological variability in space and time: insights gained from the US LTER program. BioScience 53:57–67 doi: 10.1641/0006-3568(2003)053[0057:EVISAT]2.0.CO;2

Leibold MA, Holyoak M, Mouquet N, Amarasekare P, Chase JM, Hoopes MF, Holt RD, Shurin JB, Law R, Tilman D, Loreau M, Gonzalez A (2004) The metacommunity concept: a framework for multi-scale community ecology. Ecol Lett 7:601–613 doi: 10.1111/j.1461-0248.2004.00608.x

Levin SA, Muller-Landau HC, Nathan R, Chave J (2003) The ecology and evolution of seed dispersal: a theoretical perspective. Annu Rev Ecol Evol Syst 34:575–604 doi:10.1146/annurev.ecolsys.34.011802.132428

Logue JB, Mouquet N, Peter H, Hillebrand H (2011) Empirical approaches to metacommunities: a review and comparison with theory. Trends Ecol Evol 26:482–91 doi: 10.1016/j.tree.2011.04.009

Mabry, KE (2014) Effects of sex and population density on dispersal and spatial genetic structure in brush mice. J Mammal 95:981–991 doi: 10.1644/14-MAMM-A-008

Mabry KE, Shelley EL, Davis KE, Blumstein DT, van Vuren DH (2013) Social mating system and sex-biased dispersal in mammals and birds: a phylogenetic analysis. PLoS ONE 8:1–9 doi: 10.1371/journal.pone.0057980

Matthysen E (2005) Density-dependent dispersal in birds and mammals. Ecography 28:403–416 doi: 10.1111/j.0906-7590.2005.04073.x

Meerburg BG, Singleton GR, Kijlstra A (2009) Rodent-borne diseases and their risks for public health. Crit Rev Microbiol 35:221–270 doi: 10.1080/1040841090298983

Morris DW, Diffendorfer JE (2004) Reciprocating dispersal by habitat-selecting white-footed mice. Oikos 107:549–558 doi: 10.1111/j.0030-1299.2004.12895.x

Mundt CC, Sackett KE, Wallace LD, Cowger C, Dudley JP (2009) Long-distance dispersal and accelerating waves of disease: empirical relationships. Am Nat 173:456–466 doi: 10.1086/597220.

Murray BGJ (1967) Dispersal in vertebrates. Ecology 48:975–978 doi: 10.2307/1934544

Myers P, Lundrigan BL, Hoffman SMG, Haraminac AP, Seto SH (2009) Climate-induced changes in small mammal communities of the Northern Great Lakes Region. Glob Change Biol 15:1434–1445 doi: 10.1111/j.1365-2486.2009.01846.x

Nathan R (2006) Long-distance dispersal of plants. Science 313:786–788 doi: 10.1126/science.1124975

Nathan R, Perry G, Cronin JT, Strand AE, Cain ML (2003) Methods for estimating long-distance dispersal. Oikos 103:261–273 doi: 10.1034/j.1600-0706.2003.12146.x

Pasinelli G, Walters JR (2002) Social and environmental factors affect natal dispersal and philopatry of male Red-cockaded Woodpeckers. Ecology 83:2229–2239 doi: 10.1890/0012-9658(2002)083[2229:SAEFAN]2.0.CO;2

Pérez-González J, Carranza J (2009) Female-biased dispersal under conditions of low male mating competition in a polygynous mammal. Mol Ecol 18:4617–4630 doi: 10.1111/j.1365-294X.2009.04386.x.

Rehmeier RL, Kaufman GA, Kaufman DW (2004) Long-distance movements of the deer mouse in tallgrass prairie. J Mammal 85:562–568 doi: 10.1644/1383956

Rodrigues AMM, Johnstone RA (2014) Evolution of positive and negative density-dependent dispersal. Proc R Soc Lond B Biol Sci 281:20141226–20141226 doi: 10.1098/rspb.2014.1226

Royle JA, Chandler RB, Sollmann R, Gardner B (2013) Spatial capture-recapture. Academic Press

Schloss CA, Nuñez TA, Lawler JJ (2012) Dispersal will limit ability of mammals to track climate change in the Western Hemisphere. Proc Natl Acad Sci USA 109:8606–8611 doi: 10.1073/pnas.1116791109.

Shonfield J, Do R, Brooks RJ, McAdam AG (2013) Reducing accidental shrew mortality associated with small-mammal livetrapping I: an inter- and intrastudy analysis. J Mammal 94: 745–753 doi: 10.1644/12-MAMM-A-271.1

Stamps JA (1991) The effects of conspecifics on habitat selection in teritorial species. Behav Ecol Sociobiol 28:29–36 doi: 10.1007/BF00172136

Stickel LF (1960) Peromyscus ranges at high and low population densities. J Mammal 41:433–441 doi: 10.2307/1377530

Stickel LF (1968) Home range and travels. In: King JA (eds) Biology of peromyscus (rodentia). Special publication no. 2, Stillwater, OK pp: 373–411

Taitt MJ (1981) The effect of extra food on small rodent populations: I. deermice (Peromyscus maniculatus). J Anim Ecol 50:111–124 doi: 10.2307/4035

van de Pol, M. and J. Wright (2009) A simple method for distinguishing within-versus between-subject effects using mixed models. Anim Behav 77: 753–758 doi: 10.1016/j.anbehav.2008.11.006

Van Horne B (1980) Deomgraphy of Peromyscus maniculatus populatons in seral stages of coastal coniferous forest in southeast Alaska. Can J Zool 59:1045–1061 doi: 10.1139/z81-146

Vellend M (2010) Conceptual synthesis in community ecology. Q Rev Biol 85:183–206 doi: 10.1086/652373

Wojan CM, Knapp SM, Mabry KE (2015) Spatial variation in population density affects dispersal behavior in brush mice. Ecology 96:1661–1669 doi: 10.1890/14-1661.1

Wood BA, Cao L, Dearing MD (2010) Deer mouse (Peromyscus maniculatus) home-range size and fidelity in sage-steppe habitat. West N Am Nat 70:345–354 doi: 10.3398/064.070.0307

Wood CM, McKinney ST (2015) Record long-distance movement of a deer mouse, Peromyscus maniculatus, in a New England montane boreal forest. Can Field-Nat 129:10–11 doi: 10.22621/cfn.v129i2.1699

